# SHORT-TERM EFFECTS OF CRUDE BOTANICAL EXTRACTS AND RHIZOSPHERE SOILS ON SOIL RESPIRATION AND β-GLUCOSIDASE ACTIVITY

**DOI:** 10.64898/2026.10.26.761826

**Authors:** Solomon Adejoro, Zoe Zebedee

## Abstract

Plant-derived compounds may modify microbial carbon turnover, but their effects on soil respiration and enzyme activity are poorly documented. This laboratory study compared crude ethanolic extracts and rhizosphere soils of garlic, ginger, turmeric, moringa and aloe vera with untreated soil during a 5-week incubation (n = 3). The extracts were not chemically characterized. Extracts reduced basal respiration and cumulative CO□-C relative to the control, with garlic extract giving the most persistent reduction, whereas rhizosphere soils showed stimulated CO□-C release from week 2 onward. β-Glucosidase activity was 4.3% lower under garlic extract, 14.0% lower under aloe vera extract and 57.4% lower in garlic rhizosphere soil (60.08 vs. 141.17 µg pNP g□^1^ dry soil h□^1^), whereas turmeric extract and ginger rhizosphere soil increased it. Extracts also reduced culturable bacterial, yeast and mold counts. Apparent soil organic matter was higher under several treatments but cannot be separated from added carbon. Biomass estimated from substrate-induced respiration is affected by antimicrobial compounds, so the derived quotients are exploratory. These preliminary results show treatment-specific responses; their field relevance requires characterized extracts, appropriate controls and longer studies.

## 1. Introduction

Soil holds about 1,500 Pg of organic carbon in the upper metre, more than the atmosphere (800 Pg C) and living vegetation (500 Pg C) combined (FAO and ITPS, 2015), and its persistence depends largely on heterotrophic microbial decomposition (Paul, 2014). Extracellular β-glucosidase (EC 3.2.1.21) hydrolyzes cellobiose to glucose, a rate-limiting step in cellulose breakdown, and is widely used as an indicator of soil biological quality (Esen, 1993; Tabatabai, 1994; Adetunji et al., 2017). Medicinal plants produce organosulfur compounds, polyphenols, flavonoids, alkaloids and essential oils with antimicrobial and enzyme-inhibitory properties (Rachel, 2013), yet their influence on soil carbon turnover is poorly characterized. Living roots add a different input: exudates of sugars, amino acids and organic acids that alter microbial activity and enzyme expression differently from pulsed applications of plant extracts (Bais et al., 2006). Eco-physiological quotients such as the metabolic quotient (qCO□, basal respiration per unit of microbial biomass carbon) are used to interpret microbial responses to such inputs (Anderson and Domsch, 1990), but their interpretation is difficult when the inputs are themselves antimicrobial.

This study compared crude ethanolic extracts and rhizosphere soils of garlic (Allium sativum), ginger (Zingiber officinale), turmeric (Curcuma longa), moringa (Moringa oleifera) and aloe vera (Aloe vera) in a 5-week laboratory incubation. We hypothesized that (1) extracts would suppress basal respiration and β-glucosidase activity in a species-specific manner, garlic most strongly; (2) rhizosphere soils would respond differently from extract treatments; and (3) indices derived from substrate-induced respiration would differ among treatments but, because that method is affected by antimicrobial compounds, would be of exploratory value only.

## 2. Materials and Methods

### 2.1 Experimental Site and Soil Sampling

Surface soil samples (0–15 cm depth) were collected from the Teaching and Research Farm of the Federal University of Technology, Akure (FUTA), Ondo State, Nigeria. The site is situated within the tropical rainforest agro-ecological zone of southwestern Nigeria. Collected bulk field soils were air-dried at ambient room temperature, thoroughly homogenized, and passed through a 2-mm stainless steel sieve to remove coarse organic debris, plant roots, and gravel. Replicate plastic pots were filled with 10 kg of prepared soil to support plant cultivation for rhizosphere soil generation.

### 2.2 Plant Cultivation, Crude Extract Preparation, and Treatment Administration

Propagules of garlic (Allium sativum), ginger (Zingiber officinale), turmeric (Curcuma longa), moringa (Moringa oleifera), and aloe vera (Aloe vera) were established in 10-kg soil pots under screenhouse conditions. Plants were maintained for three months with standardized bi-weekly watering regimes to allow the establishment of active, conditioned rhizosphere environments.

For crude extract preparation, fresh vegetative and root tissues from the five plant species were harvested, washed thoroughly under running tap water to remove adhering soil particles, chopped into uniform fragments, and mechanically macerated. Plant tissues were soaked in 70% ethanol for 72 hours to extract soluble bioactive constituents. The resulting suspensions were filtered through multi-layered muslin cloth, and the ethanol solvent was removed via gentle heating in a warm water bath to yield concentrated crude aqueous extracts (extraction ratio and yield are not available). It is explicitly noted that these preparations represent crude, unstandardized botanical fractions rather than isolated, purified chemical standards; chromatographic profiling (such as HPLC or GC-MS) was not performed on the crude fractions.

Extract characteristics and carbon load. No chromatographic profiling was performed. Extract carbon, pH, total phenolics and residual ethanol were not recorded, so the carbon added per jar is unknown (an upper-bound estimate is given in Section 4.4). Treatment effects are attributed to the crude extract matrix as a whole, not to any individual compound.

For the laboratory incubation experiment, 400 g of air-dried, sieved soil was weighed into sealed plastic incubation jars. Crude extracts were administered at a standardized application volume of 10 mL per jar, and soil moisture was adjusted to 60% of water-holding capacity. Authentic rhizosphere soil was collected by carefully uprooting the 3-month-old screenhouse plants, gently shaking non-adhering bulk soil from the root systems, recovering the closely adhering rhizosphere soil, and passing it through a 2-mm sieve before jar placement. Untreated control jars received an equivalent volume (10 mL) of distilled water without botanical amendment. No ethanol-evaporated solvent control and no carbon-equivalent control (e.g. a glucose solution matched to the extract carbon load) were included, so effects of the extract carrier and of added carbon cannot be separated from those of secondary metabolites. The history of the control soil relative to the pot-grown soils is not documented, so the rhizosphere–control contrast may be confounded by pot ageing. The experimental unit was one incubation jar; each treatment had three jars, and respiration was measured on the same jars each week. All treatment jars were incubated in continuous darkness at ambient laboratory temperature (25–28 °C) for a duration of 5 weeks.

### 2.3 Measurement of Soil Basal and Substrate-Induced Respiration

Soil basal respiration (SR) was measured weekly throughout the 5-week incubation period using the static alkali entrapment method (Anderson, 1982; Stotzky, 1965). Glass beakers containing 10 mL of 0.5 M sodium hydroxide (NaOH) solution were suspended inside the sealed incubation jars directly above the soil surface to capture evolved carbon dioxide (CO□-C). Concurrently, empty blank jars containing NaOH trapping solution without soil were incubated under identical conditions to quantify ambient background atmospheric carbon dioxide. Trapping solutions were harvested and replaced at 7-day intervals.

At the conclusion of each weekly interval, 1.0 M barium chloride (BaCl□) solution was added to the harvested alkali to precipitate absorbed carbonate ions as insoluble barium carbonate (BaCO□). The remaining unneutralized NaOH was back-titrated against standardized 0.5 M hydrochloric acid (HCl) using a phenolphthalein indicator. The mass of evolved CO□-C was calculated based on stoichiometric consumption. For substrate-induced respiration (SIR), soil subsamples were amended with analytical-grade D-glucose at an application rate of 4.0 mg glucose per gram of dry soil, and evolved CO□ was captured over an initial 6-hour incubation window.

### 2.4 Spectrophotometric Assay of β-Glucosidase Activity

Soil β-glucosidase activity was determined using the colorimetric method of Tabatabai (1994). Subsamples of 1.0 g of air-dried, sieved soil (< 2 mm) were placed into 50-mL Erlenmeyer flasks and treated sequentially with 0.25 mL of analytical-grade toluene, 4.0 mL of Modified Universal Buffer (MUB; adjusted to pH 6.0), and 1.0 mL of 0.05 M p-nitrophenyl-β-D-glucopyranoside (PNG) substrate solution. Flasks were sealed and incubated at 37 °C for exactly 1 hour.

Enzymatic hydrolysis was halted by adding 1.0 mL of 0.5 M calcium chloride (CaCl□) and 4.0 mL of 0.1 M tris(hydroxymethyl)aminomethane (THAM) buffer adjusted to pH 12.0. Soil suspensions were filtered through Whatman No. 2 filter paper, and the yellow coloration produced by the released p-nitrophenol (pNP) was quantified spectrophotometrically at 410 nm against a pNP analytical calibration curve. Soil controls without substrate addition during incubation were analyzed simultaneously to correct for background soil optical absorbance. Enzyme activity was expressed as micrograms of p-nitrophenol released per gram of dry soil per hour (μg pNP g□^1^ dwt h□^1^).

### 2.5 Soil Chemical Properties and Organic Matter Determination

Soil pH was determined in a 1:2 (weight/volume) soil-to-distilled-water suspension using a calibrated digital glass-electrode pH meter. Soil organic carbon (SOC) was quantified by the wet-oxidation Walkley-Black potassium dichromate digestion procedure (Ibitoye, 2006). Total soil organic matter (SOM) percentage was derived using the conventional van Bemmelen conversion factor:

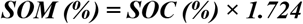

Because Walkley-Black chromic acid digestion oxidizes all readily accessible organic carbon fractions, this metric aggregates native soil carbon pools with any exogenous carbon introduced via the applied crude plant extracts. Dichromate oxidation is also sensitive to other reductants, including residual ethanol and reduced sulfur compounds such as those in Allium extracts, which would be read as additional carbon. The SOM values are therefore treated as apparent values and are not used to draw conclusions about carbon retention.

### 2.6 Enumeration of Culturable Soil Microbial Populations

Viable populations of culturable soil bacteria, yeasts, and molds were enumerated using serial dilution and the standard pour-plate technique. Aliquots of soil suspensions were inoculated onto Nutrient Agar (NA) for total bacterial enumeration (incubated at 37 °C for 24 hours) and Potato Dextrose Agar (PDA) supplemented with antibiotics for fungal (yeast and mold) enumeration (incubated at 27 °C for 48 hours). Colony-forming units were enumerated using a digital colony counter and expressed as CFU × 10□ g□^1^ dry soil.

### 2.7 Eco-Physiological Quotients and Statistical Modeling

A SIR-derived microbial biomass estimate (hereafter MBC-SIR) was calculated from substrate-induced respiration using the conversion equation of Anderson and Domsch (1978). MBC-SIR was not validated against an independent biomass measurement such as chloroform fumigation–extraction:

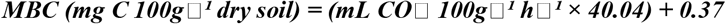

Derived eco-physiological quotients were calculated according to standard biogeochemical formulations (Anderson and Domsch, 1990; Pinzari et al., 1999). Because MBC-SIR is itself reduced by the treatments, qCO□ and qC are reported as apparent, exploratory indices:

- Microbial metabolic quotient (qCO□): The ratio of hourly basal respiration rate to microbial biomass carbon (mg CO□-C g□^1^ MBC h□^1^), indexing maintenance energy requirements.
- Carbon mineralization quotient (qM): The percentage of total soil organic carbon mineralized as carbon dioxide over the incubation period (ΣCO□-C / SOC × 100).
- Microbial carbon ratio (Cmin:Corg): The weekly ratio of mineralized carbon relative to organic carbon pools.
- Microbial carbon change rate (qC): The proportional change in MBC-SIR relative to initial conditions ((MBC□ - MBC□) / MBC□), where MBC□ is the initial MBC-SIR value.

**Methodological and Statistical Framework**

The Anderson and Domsch model assumes an uninhibited respiratory response to saturated glucose addition. In the presence of antimicrobial phytochemicals that suppress respiratory enzymes, reductions in glucose-induced CO□ efflux reflect both true reductions in physical cell biomass and metabolic enzyme inhibition. Derived quotients (qCO□ and qC) must therefore be interpreted as physiological response indicators rather than pure structural biomass ratios. Respiration data were collected repeatedly from the same jars, and the statistical treatment of this is described in Section 2.8.

**2.8 Statistical Analysis**

The extract experiment and the rhizosphere experiment were analyzed separately, each against the untreated control, in a completely randomized design with three replicate jars per treatment. Both were compared with one control, and no formal statistical comparison was made between the extract-treated and rhizosphere systems, because they differ in how the soil was prepared, in the age of the soil, and in the type of carbon input.

Respiration (SR, SIR) and cumulative CO□-C were measured weekly on the same jars. Each week was analyzed by one-way ANOVA, and treatment means were separated by Tukey’s HSD test at p ≤ 0.05 using Minitab statistical software. Variables measured once per jar (β-glucosidase activity, pH, SOM and plate counts) were analyzed in the same way, with colony counts compared on the original scale. This weekly analysis treats the weeks as independent and does not model the correlation among repeated measurements on the same jar; a repeated-measures mixed model (Treatment, Week and Treatment × Week, with jar as a random effect) would be the appropriate analysis, so the weekly significance statements are descriptive and should be read with that limitation in mind. With n = 3 jars per treatment, power to detect small differences is limited.

## 3. Results

### 3.1 Soil Basal Respiration and Substrate-Induced Respiration Kinetics

Crude botanical extracts and authentic rhizosphere soils altered soil respiratory activity in contrasting ways throughout the 5-week incubation period. Across the five extracts, weekly basal respiration (SR) and cumulative CO□-C efflux were lower than in the untreated control soil (weekly one-way ANOVA, Tukey HSD, p ≤ 0.05). Garlic extract exerted the most powerful, sustained inhibition, maintaining basal respiration rates below control levels across the entire incubation duration.

Under substrate-induced respiration (SIR) testing, extract treatments similarly suppressed glucose-triggered carbon dioxide evolution relative to untreated soil. In contrast, authentic rhizosphere soils displayed a distinct biphasic respiratory trajectory: an initial suppression during week 1, followed by respiratory stimulation from week 2 through week 5 (weekly one-way ANOVA, p ≤ 0.05). Among rhizosphere soils, moringa rhizosphere generated the highest sustained cumulative carbon dioxide evolution by the conclusion of week 5.

### 3.2 Differential Responses of Soil β-Glucosidase Activity

Extracellular β-glucosidase activity responded with pronounced divergence depending on whether botanical amendments were supplied as crude liquid extracts or as conditioned rhizosphere soil systems.

In extract-treated soils, garlic extract produced a modest but statistically significant 4.3% reduction in β-glucosidase activity, recording 135.05 μg pNP g□^1^ dwt h□^1^ compared to 141.16 μg pNP g□^1^ dwt h□^1^ in the untreated control. Aloe vera extract induced the strongest enzymatic suppression among all extracts, reducing activity by 14.0% to 121.33 μg pNP g□^1^ dwt h□^1^. Conversely, extracts of turmeric (151.33 μg pNP g□^1^ dwt h□^1^), moringa (145.26 μg pNP g□^1^ dwt h□^1^), and ginger (144.46 μg pNP g□^1^ dwt h□^1^) significantly stimulated β-glucosidase activity above control values.

In authentic rhizosphere soils, enzymatic repression was substantially amplified in specific species. Garlic rhizosphere soil caused severe enzyme inhibition, suppressing activity to 60.08 μg pNP g□^1^ dwt h□^1^—a 57.4% decrease relative to control soil (141.17 μg pNP g□^1^ dwt h□^1^). Moringa rhizosphere soil also suppressed enzyme expression (112.55 μg pNP g□^1^ dwt h□^1^), whereas ginger (153.54 μg pNP g□^1^ dwt h□^1^), aloe vera (148.00 μg pNP g□^1^ dwt h□^1^), and turmeric (147.09 μg pNP g□^1^ dwt h□^1^) rhizosphere soils exhibited significant enzymatic upregulation.

**Table 1 Overview:** The comparative treatment means for β-glucosidase activity across extracts and rhizosphere soils are detailed in Table 1a and Table 1b.

**Table 1a.** Main effects of crude botanical extracts on soil β-glucosidase activity.

| Treatment | β-Glucosidase Activity ($\mu\text{g pNP g}^{-1} \text{dwt h}^{-1}$) | Response Relative to Control (%) |
| --- | --- | --- |
| Ginger Extract | 144.46 c | +2.34% |
| Garlic Extract | 135.05 e | -4.33% |
| Turmeric Extract | 151.33 a | +7.20% |
| Moringa Extract | 145.26 b | +2.90% |
| Aloe vera Extract | 121.33 f | -14.05% |
| Untreated Control | 141.16 d | — |

\*Means within the column followed by different letters differ significantly according to Tukey's HSD test ( $p \leq 0.05$ ).

**Table 1b.** Effects of authentic plant rhizosphere soils on soil β-glucosidase activity.

| Treatment | β-Glucosidase Activity ($\mu\text{g pNP g}^{-1} \text{dwt h}^{-1}$) | Response Relative to Control (%) |
| --- | --- | --- |
| Ginger Rhizosphere | 153.54 a | +8.76% |
| Garlic Rhizosphere | 60.08 e | -57.44% |
| Turmeric Rhizosphere | 147.09 b | +4.19% |
| Moringa Rhizosphere | 112.55 d | -20.27% |
| Aloe vera Rhizosphere | 148.00 b | +4.84% |
| Control Soil | 141.17 c | — |

\*Means within the column followed by different letters differ significantly according to Tukey's HSD test ( $p \leq 0.05$ ).

**Table 2a.** Effects of crude botanical extracts on soil pH and apparent Soil Organic Matter (SOM).

| Treatment | Soil pH (1:2 H<sub>2</sub>O) | Apparent Soil Organic Matter (%) |
| --- | --- | --- |
| Ginger Extract | 5.73 ab | 13.40 b |
| Garlic Extract | 5.43 b | 19.83 a |
| Turmeric Extract | 5.40 b | 14.43 b |
| Moringa Extract | 5.73 ab | 9.17 c |
| Aloe vera Extract | 5.87 a | 14.60 b |
| Untreated Control | 5.43 b | 10.30 c |

\*Means within a column followed by different letters differ significantly according to Tukey's HSD test ( $p \leq 0.05$ ).

**Table 2b.** Effects of authentic plant rhizosphere soils on soil pH and apparent Soil Organic Matter (SOM).

| Treatment | Soil pH (1:2 H<sub>2</sub>O) | Apparent Soil Organic Matter (%) |
| --- | --- | --- |
| Ginger Rhizosphere | 6.17 bc | 15.17 a |
| Garlic Rhizosphere | 7.07 a | 14.83 a |
| Turmeric Rhizosphere | 5.60 cd | 12.70 b |
| Moringa Rhizosphere | 6.43 ab | 12.67 b |
| Aloe vera Rhizosphere | 5.17 d | 11.07 c |
| Control Soil | 5.43 cd | 10.30 c |

\*Means within a column followed by different letters differ significantly according to Tukey's HSD test ( $p \leq 0.05$ ).

### 3.3 Alterations in Soil pH and Apparent Soil Organic Matter

Soil pH and apparent soil organic matter percentages exhibited significant shifts under both extract amendments and living root conditioning.

Among crude extracts, only aloe vera significantly elevated soil pH (5.87) above the untreated control (5.43), whereas ginger and moringa extracts (5.73) showed marginal, non-significant increases, and garlic (5.43) and turmeric (5.40) remained equivalent to the control. In rhizosphere systems, garlic rhizosphere soil induced substantial alkalization, shifting pH to 7.07, while moringa rhizosphere reached 6.43. Aloe vera rhizosphere soil was slightly acidified (5.17) relative to the control.

Apparent soil organic matter (SOM) determined by Walkley-Black digestion was markedly higher in extract-treated soils. Garlic extract exhibited the highest apparent SOM at 19.83%, compared to 10.30% in the untreated control. Aloe vera (14.60%), turmeric (14.43%), and ginger (13.40%) extracts also yielded significantly elevated SOM levels, whereas moringa extract (9.17%) did not differ from the control. In rhizosphere soils, ginger (15.17%), garlic (14.83%), turmeric (12.70%), and moringa (12.67%) maintained significantly higher SOM than the unplanted control soil (10.30%).

### 3.4 Suppression of Culturable Soil Microbial Populations

Application of crude botanical extracts caused significant broad-spectrum suppression of culturable bacteria, yeasts, and molds relative to the untreated control soil (p ≤ 0.05). Ginger extract exhibited the strongest antibacterial suppression, reducing viable bacterial counts to 10.91 × 10□ CFU g□^1^ compared to 26.73 × 10□ CFU g□^1^ in the control. Yeast populations were suppressed most effectively by turmeric (5.00 × 10□ CFU g□^1^), moringa (6.00 × 10□ CFU g□^1^), and ginger (6.29 × 10□ CFU g□^1^) extracts, while mold counts were lowest under moringa (12.17 × 10□ CFU g□^1^) and aloe vera (12.49 × 10□ CFU g□^1^).

In rhizosphere soils, microbial population dynamics were more balanced. Culturable bacterial populations in ginger (23.30 × 10□ CFU g□^1^) and moringa (21.59 × 10□ CFU g□^1^) rhizosphere soils did not differ significantly from control soil, whereas garlic, turmeric, and aloe vera rhizosphere soils exhibited lower bacterial counts.

### 3.5 Trajectories of Microbial Eco-Physiological Quotients

The microbial metabolic quotient (qCO□) exhibited dynamic temporal restructuring over the 5-week incubation. During weeks 1 through 3, qCO□ decreased across extract and rhizosphere treatments relative to initial baseline levels. However, during weeks 4 and 5, qCO□ values rose across all treatments. The carbon mineralization quotient (qM) and the microbial carbon ratio (Cmin:Corg) were consistently higher in authentic rhizosphere soils than in extract-amended soils throughout the trial. Furthermore, the microbial carbon change rate (qC) turned negative at day 28 in several extract treatments, indicating an apparent decline in glucose-responsive respiration, which cannot be equated with a decline in biomass (Section 4.3). Weekly values by treatment are itemized in Supplementary Sections S3 and S4.

## 4. Discussion

### 4.1 Phytochemical Regulation of Basal Respiration and Carbon Efflux

The marked suppression of soil basal respiration by crude botanical extracts aligns with documented antimicrobial properties of medicinal plant secondary metabolites. Garlic extract exerted the most persistent and durable inhibition of carbon dioxide efflux across the 5-week trial. In botanical literature, the bioactivity of Allium species is largely driven by organosulfur constituents, notably allicin (diallyl thiosulfinate), diallyl disulfide, and diallyl trisulfide. These compounds interact with sulfhydryl (–SH) functional groups on microbial metabolic enzymes, inducing protein oxidation and enzymatic inactivation. However, because specific phytochemical constituents were not quantified or standardized via HPLC/GC-MS in the crude extracts tested here, attributing respiratory suppression exclusively to specific organosulfur fractions remains a working hypothesis rather than an analytically confirmed mechanism.

The respiratory trajectories in authentic rhizosphere soils contrasted sharply with the extract-amended systems. Following an initial week-1 lag, rhizosphere soils sustained significantly elevated carbon dioxide evolution. This stimulation is consistent with a role for continuous rhizodeposition (Hutsch et al., 2002), wherein living root systems release low-molecular-weight exudates, including glucose, fructose, organic acids, and amino acids that prime heterotrophic microbial activity and alleviate catabolic carbon starvation.

### 4.2 Differential Modulation of β-Glucosidase Activity

Extracellular β-glucosidase catalyzes the rate-limiting hydrolytic conversion of cellobiose into glucose, functioning as a critical bio-indicator of carbon turnover. In our incubation, crude extracts produced divergent enzymatic responses: garlic and aloe vera extracts moderately inhibited activity (4.3% and 14.0% reductions, respectively), whereas turmeric, moringa, and ginger extracts stimulated activity by 2.3% to 7.2%.

Enzymatic suppression was drastically amplified in garlic rhizosphere soil, where β-glucosidase activity fell by 57.4% relative to unplanted control soil. This strong inhibition could reflect continuous exposure to living garlic roots, but the rhizosphere and extract systems differ in several other respects (Section 4.4), so the contrast cannot be attributed to exposure duration alone. Organosulfur compounds and polyphenolic secondary metabolites can inactivate extracellular enzymes either by binding competitively to catalytic active sites or by altering secondary and tertiary protein conformations.

Nevertheless, reduced enzymatic expression cannot be viewed strictly as direct catalytic inhibition. Because extract applications significantly suppressed culturable bacterial and fungal populations (Table 3a), diminished β-glucosidase activity in garlic and aloe vera soils may simply reflect reduced microbial synthesis and release of extracellular enzymes rather than active-site inhibition. Conversely, the modest stimulation observed under turmeric and ginger amendments may indicate that certain phenolic constituents serve as energetic co-substrates, inducing microbial enzyme expression in less sensitive microbial taxa. Establishing the exact mechanism will require purified enzyme kinetic assays (Vmax and Km) in future investigations.

**Table 3a.**
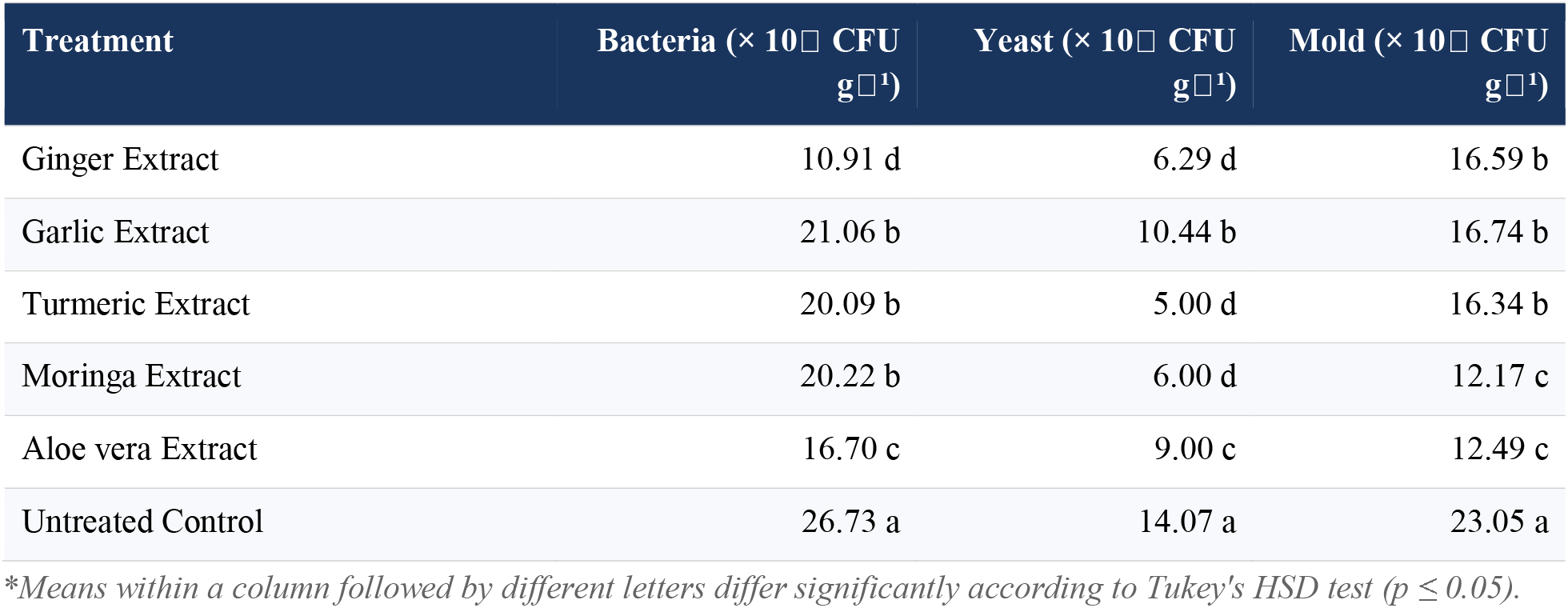
Culturable microbial populations under crude botanical extract amendments.

**Table 3b.** Culturable microbial populations in authentic plant rhizosphere soils.

| Treatment | Bacteria ($\times 10^4$ CFU g<sup>-1</sup>) | Yeast ($\times 10^4$ CFU g<sup>-1</sup>) | Mold ($\times 10^4$ CFU g<sup>-1</sup>) |
| --- | --- | --- | --- |
| Ginger Rhizosphere | 23.30 a | 6.26 d | 16.83 b |
| Garlic Rhizosphere | 16.69 c | 10.52 bc | 9.60 c |
| Turmeric Rhizosphere | 18.81 b | 10.44 c | 16.74 b |
| Moringa Rhizosphere | 21.59 a | 18.58 a | 16.95 b |
| Aloe vera Rhizosphere | 9.91 d | 9.00 cd | 22.56 a |
| Control Soil | 26.73 a | 14.07 a | 23.05 a |

\*Means within a column followed by different letters differ significantly according to Tukey's HSD test ( $p \leq 0.05$ ).

### 4.3 Limits of SIR-Derived Biomass and Eco-Physiological Quotients

A central limitation of this study concerns the estimation of microbial biomass carbon (MBC) from substrate-induced respiration (SIR) and its subsequent propagation into derived eco-physiological quotients (qCO□, qM, and qC).

The classical physiological model of Anderson and Domsch relies on measuring maximum initial respiratory response following glucose saturation, operating on the foundational assumption that respiration rate is linearly proportional to viable microbial biomass. Under typical unamended or organically fertilized soil conditions, this relationship holds robustly. However, when soils receive botanical amendments containing potent biocidal or enzyme-inhibiting metabolites, glucose uptake and catabolic respiration are directly repressed. Consequently, a lower SIR response does not necessarily indicate a smaller physical microbial biomass; it may reflect metabolic sedation, membrane disruption, or respiratory chain inhibition in viable cells.

This analytical constraint directly impacts derived quotients:

- Metabolic Quotient (qCO□ = SR / MBC): Because both basal respiration (numerator) and SIR-derived MBC (denominator) are simultaneously repressed by antimicrobial extracts, qCO□ represents a ratio of two mutually confounded catabolic metrics. The observed decline in qCO□ during weeks 1–3, followed by a sharp increase during weeks 4–5, is compatible with an initial phase of reduced catabolic activity followed by higher respiration per unit of glucose-responsive biomass, but, because both terms of the ratio are affected by the treatments, the data cannot show whether this reflects maintenance costs, detoxification, changes in community composition, or the depletion of added carbon.
- Microbial Carbon Change Rate (qC): The negative qC observed at day 28 reflects a sharp contraction in glucose-inducible respiration. This metric cannot distinguish a loss of biomass from catabolic exhaustion, dormancy or enzyme inhibition.

To resolve these ambiguities, future studies must evaluate microbial biomass using non-catabolic, structural extraction methodologies, specifically chloroform fumigation-extraction (CFE) or phospholipid fatty acid (PLFA) biomarker profiling, which quantify physical cellular pools independently of immediate respiratory competence.

### 4.4 Disentangling Acute Extract Pulses from Living Rhizosphere Dynamics and Addressing Exogenous Carbon

A critical structural distinction in this study is the fundamental difference between the two experimental systems evaluated:

1. Acute Crude Extract Amendments: Bare soil incubated with a single liquid pulse of concentrated plant extract across a short 5-week window.
2. Authentic Rhizosphere Soils: Soil dynamically conditioned over three months by living root architecture, continuous nutrient uptake, sloughed root cap cells, and fluctuating exudate gradients.

Treating these two systems as direct analytical equivalents overlooks their contrasting ecological mechanisms. The 5-week incubation trial captures immediate, transient responses to a biochemical disturbance pulse, whereas the rhizosphere soils reflect an adapted biological zone characterized by higher carbon turnover (qM and Cmin:Corg) and active microbial renewal.

Furthermore, apparent soil organic matter percentages under extract treatments (reaching 19.83% under garlic extract versus 10.30% in control) must be interpreted with extreme caution. Standard Walkley-Black wet-oxidation digests all oxidizable organic carbon present in the sample. Applying 10 mL of crude plant extract introduces an exogenous load of plant-derived sugars, organic acids, and suspended colloidal carbon into the incubation jars. Therefore, the observed elevation in SOM cannot be ascribed entirely to retarded mineralization of native soil humus. Without a complete carbon mass balance, quantifying baseline soil carbon, total carbon added via extracts, cumulative gaseous CO□-C evolved, and residual organic carbon fractions, claims that botanical extracts ‘conserve’ soil carbon remain unverified.

The size of the reported SOM difference also needs scrutiny. Taking the control value of 10.30% SOM (5.97% SOC) and the garlic-extract value of 19.83% SOM (11.50% SOC), and assuming these values are expressed on a dry-soil basis, the difference corresponds to roughly 22 g of carbon in 400 g of soil. A 10 mL aliquot would have to contain more than 2 g C mL□^1^ to supply this, whereas even a saturated sucrose solution contains less than 0.5 g C mL□^1^. Added extract carbon therefore cannot account for the difference on its own. Other possible contributors are interference with the dichromate digestion by reduced compounds in the extract (for example organosulfur compounds in garlic or residual ethanol), incomplete homogenization of the extract-treated soil, or an error in the analytical data. The same applies, to a lesser degree, to the rhizosphere soils, where root fragments passing the 2-mm sieve would also be digested. The SOM values are therefore treated as apparent values and are not used to draw conclusions about carbon retention.

### 4.5 Agronomic Considerations, Ecological Trade-Offs, and Research Priorities

The suppression of carbon dioxide evolution and β-glucosidase activity by garlic-derived compounds demonstrates clear biological activity, but translating these laboratory observations into agronomic practice presents significant ecological trade-offs:

- Disruption of Essential Mineralization: Slowing organic matter decomposition simultaneously impedes the biological mineralization of essential plant nutrients, including nitrogen, phosphorus, and sulfur. An amendment that suppresses microbial activity to conserve carbon may inadvertently induce crop nutrient deficiency in field production systems.
- Short-Term Lab Incubation versus Field Reality: Laboratory incubations in closed jars eliminate field leaching, photolysis, aggregate protection, and microbial colonization from subsoil layers. The persistence of bioactive secondary metabolites in open soil environments is typically far shorter due to rapid microbial degradation and chemical adsorption onto mineral surfaces.

To advance beyond these preliminary findings, future research must satisfy five experimental prerequisites:

1. Phytochemical Profiling: Full chromatographic identification and quantification (HPLC/GC-MS) of marker compounds in botanical extracts.
2. Matrix Controls: Inclusion of empty solvent-evaporated and carbon-equated amendment controls to isolate active secondary metabolites from carrier and nutritional matrix effects.
3. Structural Biomass Validation: Parallel determination of microbial biomass using chloroform fumigation-extraction (CFE) and PLFA analysis alongside SIR.
4. Complete Carbon Mass Balancing: Tracing native versus exogenous carbon turnover using isotopic (^13^C) labeling techniques.
5. Enzyme Kinetics and Field Trials: Evaluating Michaelis-Menten kinetic constants (Vmax and Km) for β-glucosidase and conducting multi-season agronomic field trials.

### 4.6 Limitations

The main limitations of this study are as follows. (i) The extracts were crude, unstandardized and uncharacterized, so no response can be attributed to a particular compound, and the missing solvent and carbon-equivalent controls prevent separation of carrier and carbon effects from those of secondary metabolites. (ii) Microbial biomass was estimated only from SIR, which was itself suppressed by the treatments; qCO□ and qC are therefore exploratory, and no independent biomass measurement was made. (iii) Respiration was measured repeatedly on the same three jars per treatment; the weekly ANOVA used (Section 2.8) does not model the correlation among repeated measurements, and power is limited. (iv) The incubation lasted five weeks, used one soil, one extract application rate and a single application, and was run in closed jars at ambient laboratory temperature, so persistence of effects, dose–response and field relevance are untested. (v) The extract and rhizosphere systems differ in design (a single liquid pulse on bulk soil versus three months of root conditioning in pots, with possible root debris in the sieved soil), the history of the control soil is not documented, and the comparison between them is descriptive only. (vi) Culturable counts represent a small, medium-dependent fraction of the community and, together with lower enzyme activity, do not by themselves indicate improved or reduced soil health.

## 5. Conclusions and Recommendations

In this short laboratory incubation, crude botanical extracts and rhizosphere soils were associated with distinct, plant-specific differences in respiration, extracellular enzyme activity and culturable microbial populations:

1. Crude botanical extracts, particularly garlic (Allium sativum), reduced basal soil respiration and cumulative carbon dioxide evolution under controlled laboratory conditions, whereas rhizosphere soils showed higher carbon efflux from week 2 onward, consistent with rhizodeposition.
2. β-Glucosidase activity differed among treatments: it was much lower in garlic rhizosphere soil (>57% reduction) and higher under turmeric extract and in ginger rhizosphere soil. Whether this reflects enzyme inhibition or fewer enzyme-producing microbes was not determined.
3. Apparent soil organic matter was substantially higher in extract-amended soils, but this effect cannot be separated from the exogenous carbon introduced by the uncharacterized crude extracts.
4. Biomass estimated from substrate-induced respiration is biased by antimicrobial compounds, so the derived quotients (qCO□, qC) are exploratory and do not support conclusions about microbial carbon-use efficiency or carbon conservation.

### Recommendations

Garlic extract and garlic rhizosphere soil showed the largest responses and are candidates for further study. However, before botanical extracts or companion planting strategies can be recommended for agronomic carbon management or soil health enhancement, rigorous studies incorporating phytochemical standardization, structural biomass determinations, mass-balance carbon accounting, and multi-season field evaluations are essential.

## Acknowledgements

None. This research received no external funding

## Author contributions

Adejoro: conceptualization, supervision, methodology; Zoe Zebedee: investigation, formal analysis, writing – original draft.

